# inTRACKtive — A Web-Based Tool for Interactive Cell Tracking Visualization

**DOI:** 10.1101/2024.10.18.618998

**Authors:** Teun A.P.M. Huijben, Ashley G. Anderson, Andrew Sweet, Erin Hoops, Connor Larsen, Kyle Awayan, Jordão Bragantini, Merlin Lange, Chi-Li Chiu, Loïc A. Royer

## Abstract

We introduce inTRACKtive, an innovative web-based tool for interactive visualization and sharing of large 3D cell tracking datasets, eliminating the need for software installations or data downloads. Built with modern web technologies, inTRACKtive enables researchers to explore cell-tracking results from terabyte-scale microscopy data, conduct virtual fate-mapping experiments, and share these results via simple hyperlinks. The platform powers the Virtual Embryo Zoo, an online resource showcasing cell-tracking datasets from state-of-the-art light-sheet embryonic microscopy of six model organisms. inTRACKtive’s open-source code allows users to visualize their own data or host customized viewer instances. By providing easy access to complex tracking datasets, inTRACKtive offers a versatile, interactive, collaborative tool for developmental biology.

---

The field of developmental biology is undergoing a profound transformation, driven by groundbreaking advances across multiple disciplines.^1^ In particular, the convergence of state-of-the-art light-sheet microscopy and cutting-edge image analysis algorithms is redefining our ability to study and understand embryogenesis *in vivo*.^2^ Light-sheet microscopy enables rapid, large-scale, multicolor imaging of developing embryos in unprecedented detail, allowing scientists to capture dynamic processes as they unfold in real-time.^3–5^ Simultaneously, the advent of advanced cell tracking technologies, such as Ultrack,^6^ Linajea,^7, 8^ Elephant,^9^ TGMM,^10^ Trackmate,^11^ Mastodon^12^ and TrackAstra,^13^ has made it possible to follow individual cells throughout embryonic development with remarkable precision and speed.^14^ These innovations, which have emerged in the past two decades, are only now realizing their full potential for embryology, offering the capability to track almost all cells in an embryo, from the earliest stages of development to fully gastrulated embryos and beyond. This unprecedented level of detail opens the door to answering fundamental questions about tissue development, organ formation, and the intricate orchestration of the cell behaviors that govern the entire embryo.^15, 16^

However, the size and complexity of these microscopy datasets — often reaching tens of terabytes and containing up to tens of thousands of cells across hundreds of time points — pose significant challenges. While these imaging datasets hold the potential to reveal novel biological insights, accessing and interacting with such complex data typically demands highly specialized technical expertise and substantial computational resources. Consequently, only a limited subset of researchers can fully leverage these data, creating a barrier to broader adoption. Could a more user-friendly solution be developed that enables researchers to explore and analyze cell tracks of these datasets, regardless of their computational expertise or available hardware? Existing software tools allow visualization and analysis of cell tracking data (*e*.*g*., *napari*,^17^ TrackMate’s TrackScheme,^11^ Mastodon,^12^ and CellTracksColab^18^), However, while very powerful in their respective application domains, most of these approaches require native software, have limited 3D support, or are intended for the curation of tracks; not the interactive exploration of large cell tracking datasets.

We introduce inTRACKtive, a web-based tool designed to address this challenge by providing an intuitive platform for visualizing and interacting with large 2D and 3D cell tracking datasets. With inTRACKtive, users can seamlessly explore large tracking datasets in an interactive, three-dimensional environment (Fig. 1a-b and Supplementary Video 1). One can select specific (groups of) cells and trace their lineages through time, which we validate by tracking tailbud cells during zebrafish development (Fig. S1); recapitulating previous findings that tailbud cells end up in the tail of the zebrafish embryo, particularly in the mesoderm, neural tube and tailbud itself.^16^ The platform operates entirely in a web browser, requiring no software installation or manual data downloads, making it easily accessible to any user. In addition, users can share their exact viewing configuration - complete with view settings, zoom levels, and selected cells - simply by sharing a link. This allows for effortless collaboration and dissemination of results, when the dataset is located on a publicly accessible server.

**Fig. 1.**
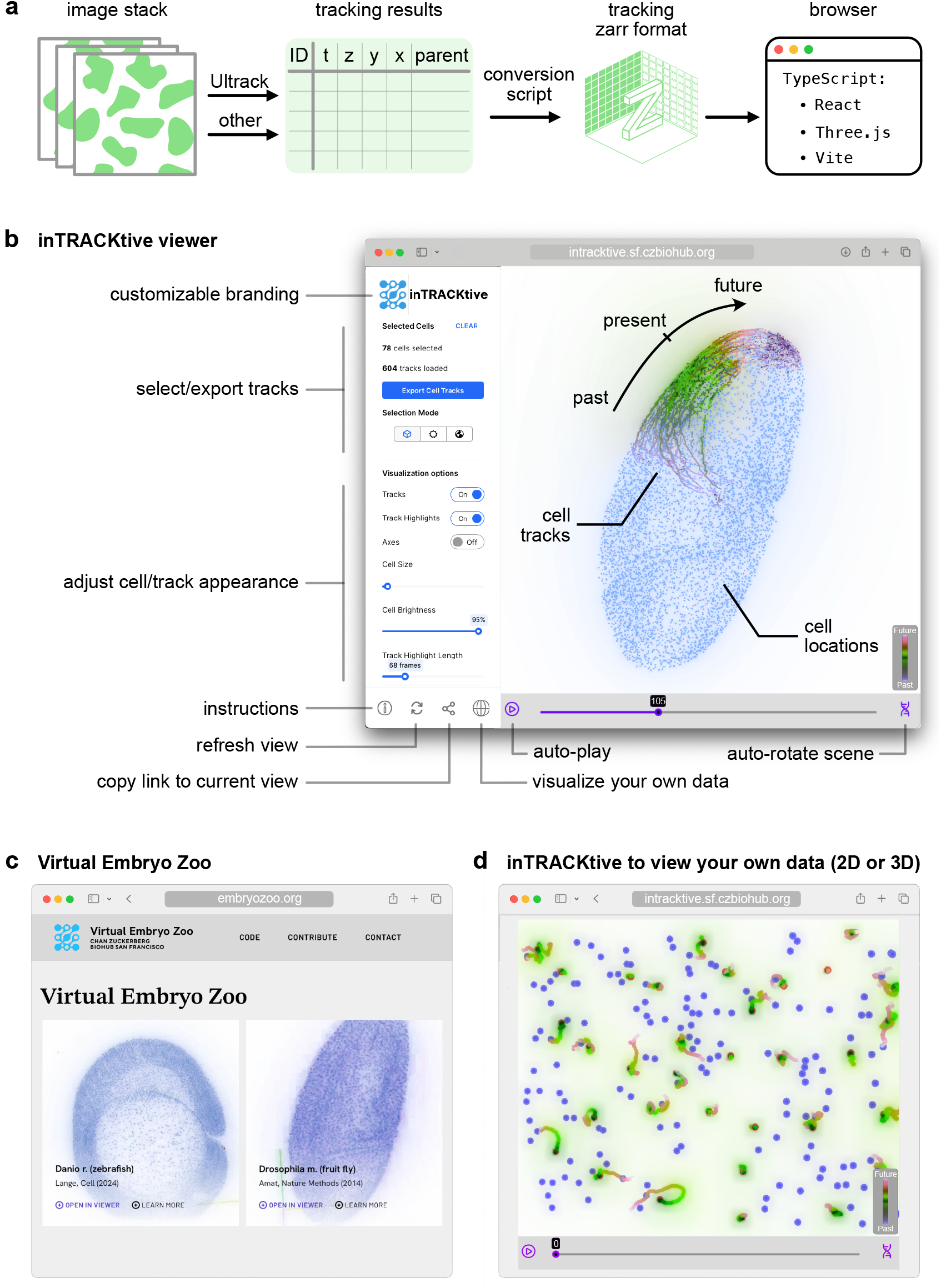
inTRACKtive cell tracking viewer and its applications. **a** inTRACKtive data pipeline. After users track their datasets with their preferred cell tracking software, the tracking results are converted to our custom Zarr format using a provided conversion script, and can be interactively visualized in the web-based tool. **b** The inTRACKtive web-viewer of cell tracking results. Users can upload their own data, select cells of interest, trace their lineages, adjust the appearance of the cells and tracks, and copy the state of the viewer into a sharable link. **c**, The Virtual Embryo Zoo encompasses six light-sheet microscopy datasets of developing embryos. The website provides a video graphical overview of the datasets, with one-click access to inTRACKtive viewer for visualizing the cell tracking data (see **b**) and the details of the data (paper, organism, authors, tracking algorithm, etc.). **d** inTRACKtive can be used to visualize one’s own 2D or 3D cell tracking data in the browser.

We used inTRACKtive to create the Virtual Embryo Zoo (embryozoo.org), an online resource that showcases the highest-quality tracked light-sheet microscopy datasets of developing embryos (Fig. 1c and Supplementary Video 2). This platform allows researchers to investigate the early embryogenesis of six widely studied model organisms (Table 1), Drosophila,^10^ zebrafish,^16^ C. elegans,^19^ ascidian,^20^ mouse,^4^ and Tribolium^21^ — via a web-based interface. Not only can users visualize these datasets, but they can also download the complete tracking data, which can be easily imported into *napari* and Python for further analysis, such as quantitative morphological analysis of track shapes, cell densities, and movement directions. In addition, the Virtual Embryo Zoo offers researchers the opportunity to contribute and showcase their own datasets (Fig. 1d), with the vision of creating a growing collaborative resource for the community.

**Table 1.**
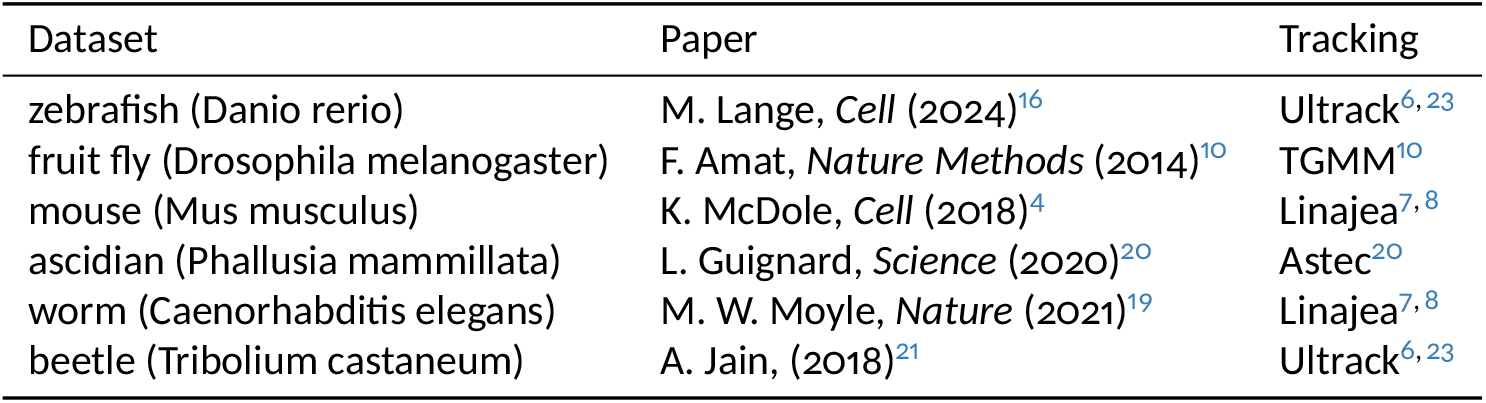
Details of the six embryo datasets portrayed in the Virtual Embryo Zoo.

inTRACKtive is an ideal tool for crafting custom celltracking visualizations. In addition to selecting and tracing individual cells, the platform allows users to color cells based on any provided attribute. For example, we demonstrate its application in Drosophila development, where cells are colored according to their velocity (Fig. S2 and Supplementary Videos 3 and 4). This functionality is not limited to velocity; it can represent any attributes of interest such as cell size, gene expression, or track length, offering extensive flexibility for exploring complex datasets.

Technically, inTRACKtive is built using modern web technologies, including TypeScript, React, and three.js, with Vite as the bundler and Vercel for deployment. The platform is optimized for performance, allowing users to interact with large-scale datasets. The app runs entirely client-side and only relies on static hosting for both the application and data. This makes it easy and inexpensive to self-host the application and make your own data accessible. One key innovation is the use of a specialized cell tracking format based on Zarr^22^ (see Fig. S3 and Supplementary Methods), which enables asynchronous, lazy data loading. This ensures that tracking results from large imaging datasets can be explored seamlessly without significant delays or computational overhead. The tool works in any browser, including tablets and smartphones, offering flexibility in how users interact with the data.

Looking ahead, there is potential to expand inTRACKtive’s capabilities. One exciting possibility is the incorporation of imaging data alongside cell-tracking results. Another area for development is further customization of track visualization and analysis of the selected tracks. While this functionality is not yet available directly in inTRACKtive, the platform allows the export of a user-selected subset of data to a `csv` format that can be further analyzed and visualized in tools like *napari*^*17*^. Thus, inTRACKtive is an ideal platform for sharing and disseminating complex cell tracking results, combining ease of use with the flexibility to integrate with more specialized software for advanced analyses.

In summary, inTRACKtive offers a powerful, versatile tool for the interactive visualization of cell tracking data, eliminating the need for complex software installations or large data downloads. Beyond visualization, inTRACKtive enables users to select individual cells, trace their lineages, and investigate both their ancestors and descendants, facilitating deeper insights into developmental processes. The platform serves multiple use cases: as a gateway for exploring datasets in the Virtual Embryo Zoo, as a tool for visualizing a researcher’s own cell tracking data (easily shared via a link), and as a customizable viewer for users who want to host their own instances. Conversion scripts for exporting cell tracking results into the Zarr format used by inTRACKtive are provided. Furthermore, we provide an intuitive command line and Python API, including a *napari* widget and a Jupyter Notebook widget, enabling users to launch inTRACKtive directly from a terminal, Jupyter Notebook or *napari* (Supplementary Video 5). This eliminates the need to spin up a local host or convert data to Zarr manually, allowing for seamless visualization of custom tracking datasets.

Importantly, inTRACKtive is freely available and opensource (github.com/royerlab/inTRACKtive). By enabling the creation of the Virtual Embryo Zoo, inTRACKtive has the potential to become a valuable resource to the cell and developmental biology community and beyond. Moreover, inTRACKtive can be used to visualize tracking data acquired with any 2D ([example](https://t.ly/K-hA7)^6^) or 3D microscopy modality from which tracking data can be obtained, from organoids down to single molecules. inTRACKtive offers a unique, browser-based playground for interacting with state-of-the-art microscopy tracking datasets.

## Supporting information

inTRACKtive web-viewer

Virtual Embryo Zoo website

Color cell labeling in inTRACKtive based on any attribute

Visualizing velocity, density, and structure tensor in inTRACKtive

Using inTRACKtive from the command line, from napari, from a Jupyter notebook, and within a Jupyter notebook

## Supplementary Videos

1. [inTRACKtive web-viewer](https://vimeo.com/1019958933/a61cfa4120);
2. [Virtual Embryo Zoo website](https://vimeo.com/1019959289/edfcb4d6a7);
3. [Color cell labeling in inTRACKtive based on any attribute](https://vimeo.com/1037262379/86bd332f4e);
4. [Visualizing velocity, density, and structure tensor in inTRACKtive](https://vimeo.com/1088515499/483f98cd9d);
5. [Using inTRACKtive from the command line, from napari, from a Jupyter notebook, and *within* a Jupyter notebook](https://vimeo.com/1088515499/483f98cd9d);

## Data availability statement

The tracking data presented in this study are available for download from the Virtual Embryo Zoo website (embryozoo.org). The datasets are either sourced from the original authors or tracked by us using the Ultrack^6, 23^ algorithm (see Table 1). All data has been converted and normalized to a common (see Fig. S3) format compatible with inTRACKtive and *napari*, ensuring seamless interaction and visualization.

## Code availability statement

The repository for inTRACKtive, including documentation, examples code, and videos, can be found at github.com/royerlab/inTRACKtive.

## Acknowledgements

We extend our gratitude to all the original authors who kindly provided data and guidance: Philipp Keller, Hari Shroff, Kate McDole, Akanksha Jain, Robert Haase, and Grégoire Malandain. We thank the Royer group at CZ Bio-hub SF and Estibaliz Gómez-de-Mariscal for feedback on inTRACKtive and the Virtual Embryo Zoo website. We appreciate Sandy Schmid for her constructive feedback on the manuscript. We thank CZI for the Science Design System ([SDS](https://github.com/chanzuckerberg/sci-components?tab=readme-ov-file)) UI component library. Chan Zuckerberg Biohub San Francisco (CZB SF) funded this work. We thank the CZB SF donors P. Chan and M. Zuckerberg for their generous support.

## Author Contributions

J.B. and L.A.R. conceived the research. A.A., A.S., E.H, C.L. and C-L.C developed the initial version of inTRACKtive. T.A.P.M.H. collected the light-sheet datasets from the original authors, converted them into the applicable formats, and further developed inTRACKtive (guided by A.A. and A.S.). K.A. and T.A.P.M.H. built the Virtual Embryo Zoo website. J.B. performed tracking of the Tribolium and zebrafish data using Ultrack. M.L. provided insight into the zebrafish tailbud migration. T.A.P.M.H. and L.A.R. wrote the paper. T.A.P.M.H. led the project. L.A.R. supervised the research. All authors contributed to editing the manuscript.

## Supplementary Methods

The cell tracking data is structured as a table with six columns: `track id`, `t`, `z`, `y`, `x`, and `parent track id`, where each row represents a cell at a single time point (Fig. S3a). The `track id` column provides the identifier for each track, while `t`, `z`, `y`, and `x` represent the time point and spatial coordinates of a specific cell (providing `z` is optional). Note that the order of the columns is not important, since they are accessed by name. The `parent track id` column specifies the `track id` of the parent track, with a value of − 1 indicating no parent. To enable fast and asynchronous loading in inTRACKtive, this data is stored in a specialized Zarr^22^ format optimized for efficient interaction. The Zarr format allows users to select a single cell at a specific time point and visualize its trajectory, as well as those of its ancestors and descendants (Fig. S3b). The cell tracking data is organized into a Zarr store comprising four Zarr arrays: `points`, `points to tracks`, `tracks to tracks`, and `tracks to points` (Fig. S3c-d). Within this context, cells are referred to as `points`. Below, we detail these four arrays and their roles before describing their storage in the Zarr format.

### Four matrices

The `points` matrix (Fig. S3c) is a dense, ragged array of 32-bit floats with dimensions (`n timepoints`, `num values per point` *×* `max points per timepoint`). The number of rows (`n timepoints`) corresponds to the total time points. Each row represents a single time point, containing the coordinates (`z`, `y`, `x`) of all observed points. Rows are padded with `-9999.9` to reach the `max points per timepoint`. Each point is assigned a unique identifier based on its time point (`t`) and index (`n`) within that time point: `point ID` = `t` *×* `max points per timepoint` + `n`. This matrix allows efficient retrieval of point coordinates for a given time frame.

The `points to tracks` matrix (Fig. S3c) maps point IDs to their associated `track id`. Its shape is (`n points`, `n tracks`). Each row represents a point, while columns denote tracks. A nonzero element (value = 1) in the matrix indicates that a specific point belongs to the corresponding track. Although values are not meaningful in this context, the column indices encode the `track id`. This matrix serves as an adjacency matrix to associate points with tracks.

The `tracks to tracks` matrix (Fig. S3c) describes relationships between tracks and has dimensions (`n tracks`, `n tracks`). This matrix allows the determination of ancestors and descendants of a track. Rows represent tracks, while columns indicate the related tracks belonging to the same lineage. All tracks present in a row will be loaded by the application when a respective track is selected. As related tracks, we consider all the ancestors and descendants of the respective track. A value of − 1 signifies a track without an ancestor, which happens when a cell was present in the first timepoint, or emerged and did not have a parent (no `parent track id`). A nonzero element in the matrix indicates a direct parent-child relationship, and the value itself is the `parent track id`. The values of the nonzero elements in the matrix are not essential to the application, since the column index already signifies the related `track id`, the value is only used to efficiently export the tracking data of the selected cells.

The `tracks to points` matrix (Fig. S3c) maps tracks to their constituent points. It has dimensions (`n tracks`, `n points`) and is essentially the transpose of the `points to tracks` matrix. Additionally, this array includes the point coordinates, which creates a redundancy, but allows for optimized queries to fetch point coordinates.

### Zarr format

The four matrices are stored as Zarr arrays in a single Zarr store (Fig. S3d). The `points` array is chunked by time, ensuring that only the data for a single time point is loaded during visualization. The remaining arrays (`points to tracks`, `tracks to tracks`, and `tracks to points`) are stored as compressed sparse row (CSR) matrices using `scipy.sparse`. The CSR format was specifically chosen for efficient row-retrieval, so each matrix is intended to be indexed by row. The CSR format encodes matrices efficiently with three arrays: `indices`, which contains the column indices of nonzero elements; `indptr`, which contains pointers to the start of rows; and `data`, which contains the values of nonzero elements. Our CSR arrays are essentially Zarr groups with minimal added metadata and 2-3 dense Zarr arrays. For `points to tracks`, only `indices` and `indptr` are stored, as the matrix values are irrelevant.

The final outline of the Zarr store (for the example zebrafish dataset) will look like this:

`tracks_bundle.zarr (`∼`550M`)

|-- points (198M)

|-- points_to_tracks (62M)

| |-- indices (61M)

| |-- indptr (1M)

|-- tracks_to_points (259M)

| |-- data (207M)

| |-- indices (50M)

| |-- indptr (2M)

|-- tracks_to_tracks (28M)

|-- data (19M)

|-- indices (7M)

|-- indptr (2M)

Note the relatively small size of the `indptr` arrays provides an opportunity for efficient caching and reduces the total number of row-wise requests when fetching tracks.

This structure includes minimal redundancy to optimize queries and reduce latency during data fetching. Despite the redundancy, the Zarr store has a size of only 46% of the original `CSV` file, since it is binary encoded.

### Fetching of tracks

To display tracks in the inTRACKtive viewer, the following steps retrieve the required coordinates (Fig. S3e). The first query to `points to tracks` identifies the `track id` of the selected point. The second query to `tracks to tracks` finds all related tracks, including ancestors and descendants. The third query to `tracks to points` retrieves the coordinates of all points in the identified tracks. These queries run asynchronously using `async`/`await` in TypeScript, ensuring efficient data retrieval.

### Adding cell size and color attributes

To enhance visualization, additional cell attributes can be incorporated. Firstly, each cell can be given a different size. A `radius` column in the input `tracks.csv` file and the `--add radius` flag in the conversion script will add cell radii to the `points` array as a fourth coordinate (`z`, `y`, `x`, `radius`), which is automatically read by inTRACKtive as the cell size in the rendering. Secondly, every cell can optionally be given a color based on provided attributes (see Fig. S2 for an example). Additional extra columns in `tracks.csv` and the `--add all attributes` flag during conversion will generate a fifth Zarr array named `attributes`. This array stores custom attributes for each cell as a dense, ragged array with dimensions (`n timepoints`, `num attributes` *×* `max points per timepoint`), and is chunked by time and attribute.

## Supplementary Figures

**Fig. S1.**
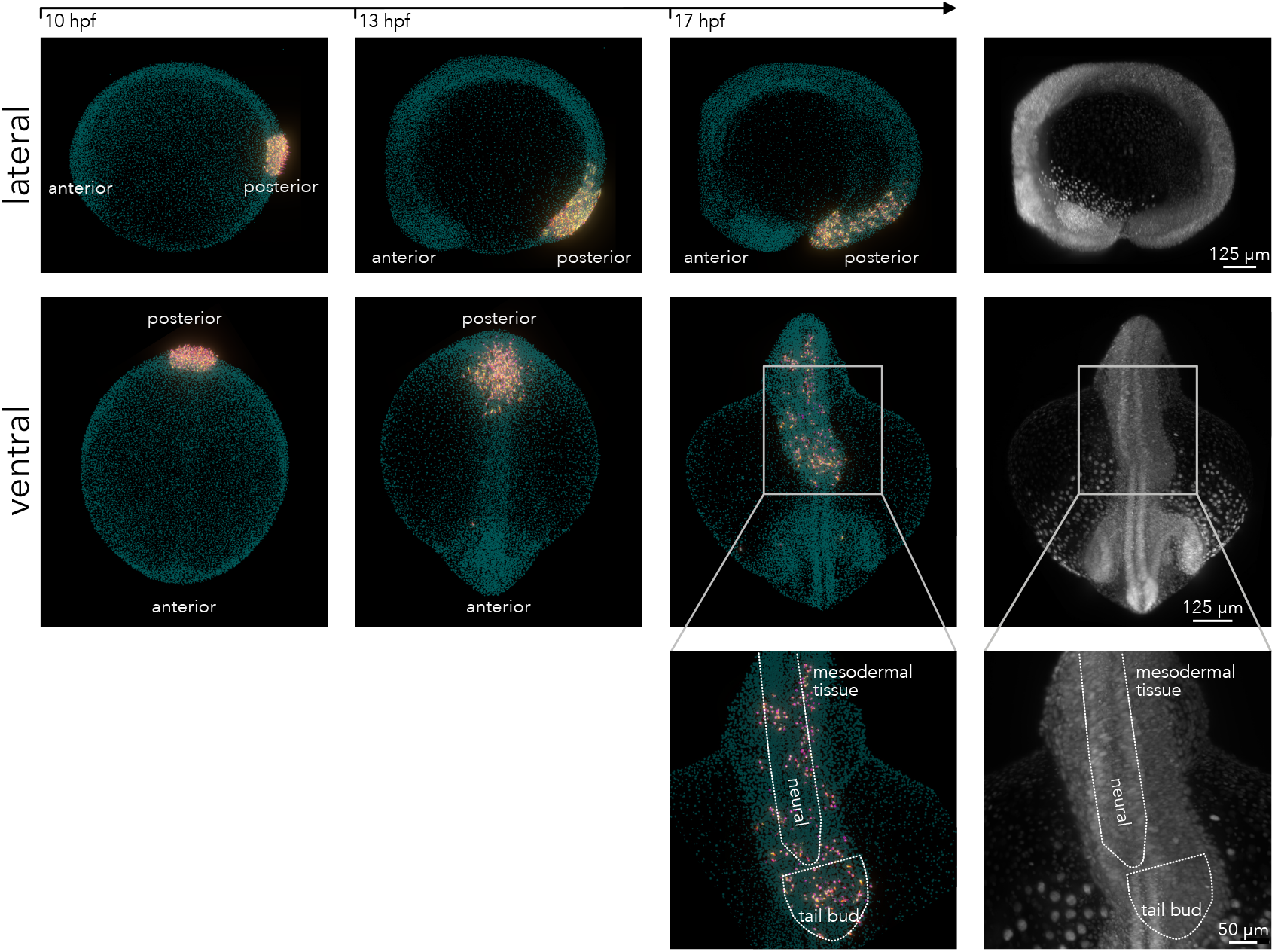
In-silico fate mapping of zebrafish tailbud cells using inTRACKtive. The fully-tracked zebrafish embryo visualized with inTRACKtive. The tailbud cells are selected within inTRACKtive at 10 hours post fertilization (hpf) and followed during development. Shown are inTRACKtive snapshots at 13 hpf (second column) and 17 hpf (third column), alongside the imaging data at 17 hpf. The zoomed panel shows that the selected tailbud cells end up in the tail of the zebrafish embryo, in particular, the mesoderm, neural tube, and tailbud itself – agreeing with previous findings.^16^

**Fig. S2.**
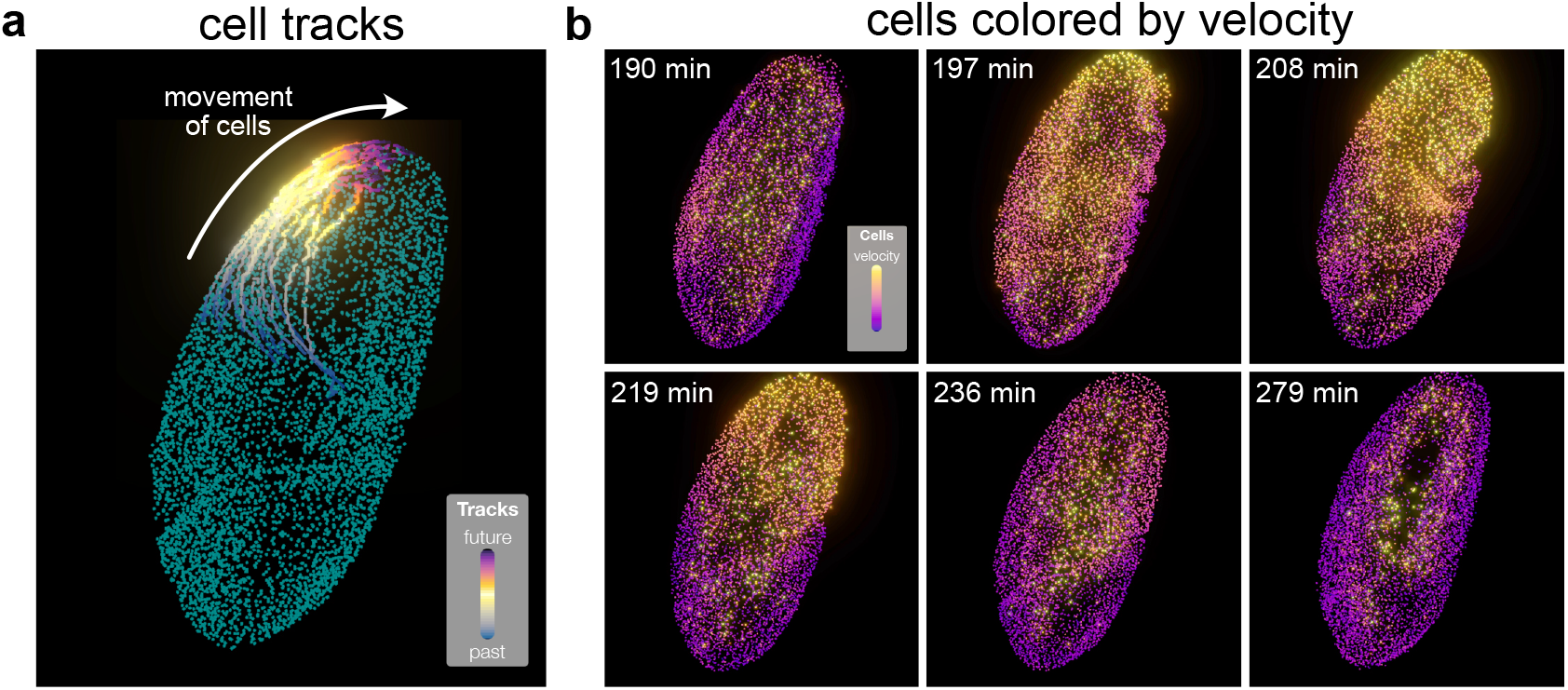
inTRACKtive allows coloring of cells based on a given attribute. **a** Fully tracked developing Drosophila melanogaster^10^ embryo, visualized in inTRACKtive. When selecting several cells, the visualized tracks show the global movement of cells during gastrulation. **b** Same embryo visualized in inTRACKtive, but with the cells colored based on their velocity. The color of the cells (yellow means high velocity) clearly displays how the wave of migration propagates through the developing embryo. Timestamps represent minutes post-fertilization.

**Fig. S3.**
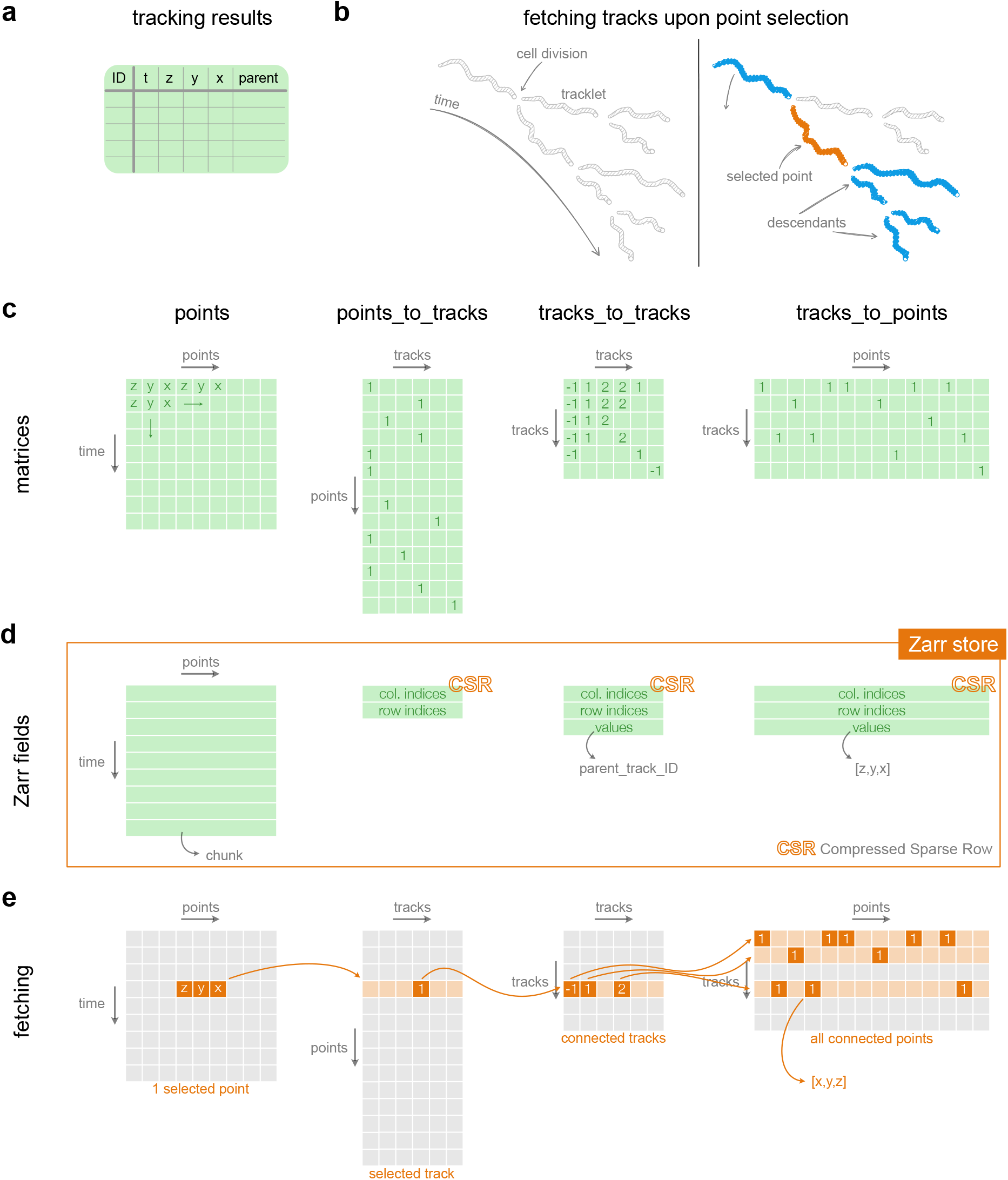
The tracking Zarr data format. **a** The start point is a table with cell tracking results. **b** The Zarr format is designed for fast and efficient fetching of tracks (ancestors and descendants) when the user selects a cell (point). **c** From the tracking results table, four matrices are constructed, one with the coordinates of all points, and three for the connections between points and tracks. **d** The four matrices of **c** are saved as Zarr arrays in one Zarr store. The points array is saved as a dense Zarr array, chunked per time. The other three are saved in the CSR (compressed sparse row) format, with fields indices (column indices), indptr (row indices), and data (values). **e** When the user selects a point, a series of three queries sequentially fetches the points of all the related tracks.

